# A Bond-Anchored Network Structure Alignment (BANSA) Method for Graphical Analysis of Protein-Protein Interfacial Regions

**DOI:** 10.1101/2023.10.16.562551

**Authors:** Justin Tam, Brian Y. Chen

## Abstract

We introduce a method to analyze and compare intermolecular bonds formed between protein-protein interactions. Utilizing the DiffBond software, we calculate potential intermolecular bonds, such as ionic bonds, hydrogen bonds, and salt bridges, based on amino acid structural and spatial parameters. This results in a graphical representation of bonds termed a bond network for each protein pair interaction. We then introduce an innovative strategy, the Bond-anchored Network Structural Alignment (BANSA), to align these networks using bond formation as anchor points. This alignment process uses the Root Mean Square Deviation (RMSD) to quantitatively assess the similarity between molecular structures. We validate the BANSA approach using several forms of analysis including a heatmap analysis, which provides a consolidated view of the entire bond network as well as a thorough comparison with existing literature. The results highlight the method’s potential to offer insights into molecular interactions across various protein pairs without a need for direct modelling of protein-protein interactions.

## I. Methods

This method aims to analyze and compare interactions between two proteins, particularly focusing on the intermolecular bonds that form between them.

Utilizing the DiffBond software, the method begins by calculating all the intermolecular bonds that could occur between two interacting proteins. Calculations for intermolecular bonds, such as ionic bonds, hydrogen bonds, and salt bridges, are based on amino acid structural and spatial parameters. This process results in a graph representation of the interfacial region called a bond network. For a comprehensive collection of protein-protein interactions, a unique bond network can be generated for each pair.

### A. Bond-anchored Network Structural Alignment (BANSA)

For structural comparison of proteins, a major obstacle is alignment of structure. The networks, generated from structural data, must also be aligned in order to perform network to network analysis. We introduce a method for aligning structures using significant bond formation as anchor points, a bond-anchored network structural alignment (BANSA).

The method requires creating k-hop neighbor groups for amino acids located on each end of a bond within the bond network. This step aggregates information about the immediate amino acid neighbors surrounding the bond ends and neighboring amino acids, providing context for each bond.

Once k-hop groups are defined, every possible combination of k-hop groups from one bond network is aligned with another in a pairwise manner. Bond Network Alignment via RMSD: For each alignment made in the previous step, the Root Mean Square Deviation (RMSD) of the full bond network is computed. RMSD is a measure to quantitatively determine the degree of similarity between two molecular structures. Lower RMSD values indicate a closer match, while higher values signify greater divergence. This exhaustive alignment process ensures that each potential correspondence between bond groups across the two networks is measured and ranked.

Finally, we filter out all the alignments that have an RMSD value below a set threshold of 2.0. The remaining alignments are then arranged in ascending order, with the lowest RMSD values indicating the most similar structures first.

This form of BANSA alignment offers a systematic way to compare and analyze the bond structures in protein-protein interactions using biochemically relevant information, specifically with bond data functioning as anchor points for alignment. BANSA provides insights into similarities and differences in molecular interactions across various protein pairs.

## II. Results

After producing aligned bond network graphs using BANSA, there are multiple scientific insights that can be extracted. Fig. 3 shows a simplified example of mutational effects on a bond network. Mutational experiments often exert diffuse bond changes in the interfacial region; we explore this property using BANSA and correlate results to known literature.

**Fig. 1.**
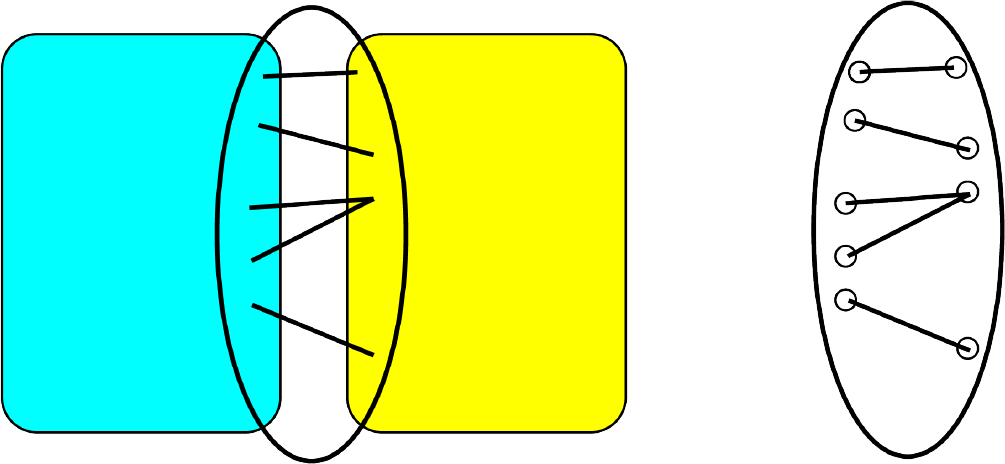
A 2D example of a bond network. The network of interactions between protein A(teal) and protein B(yellow) is extracted using DiffBond.

**Fig. 2.**
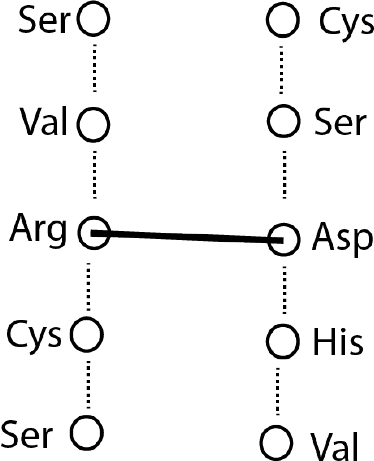
An example of a single K-hop group extracted from a network for use in structure alignment. Aggregates some neighborhood information around nodes involved in bonds.

**Fig. 3.**
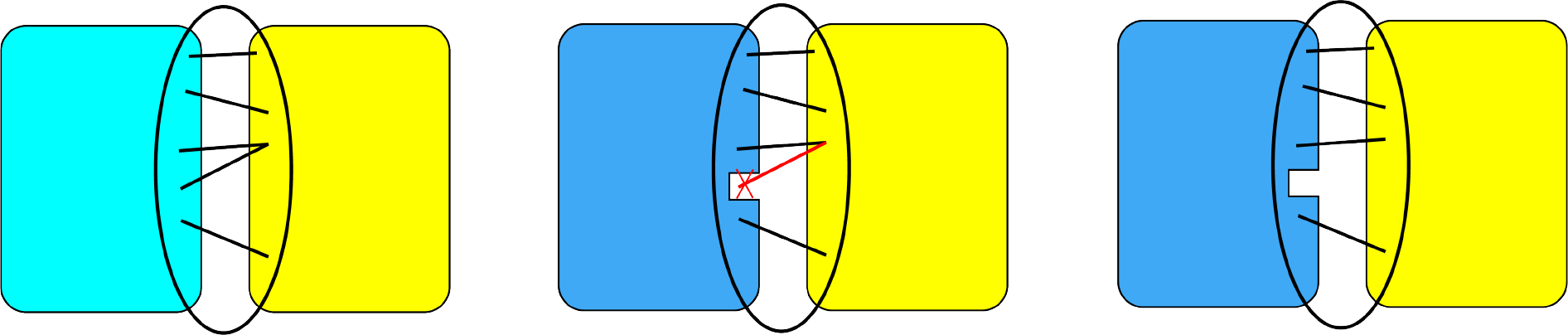
Effect of mutation in a protein on bond network. A wildtype protein (teal) compared to a mutant (blue) may have interfacial differences due to mutation which can add or remove bonds in the bond network.

**Fig. 4.**
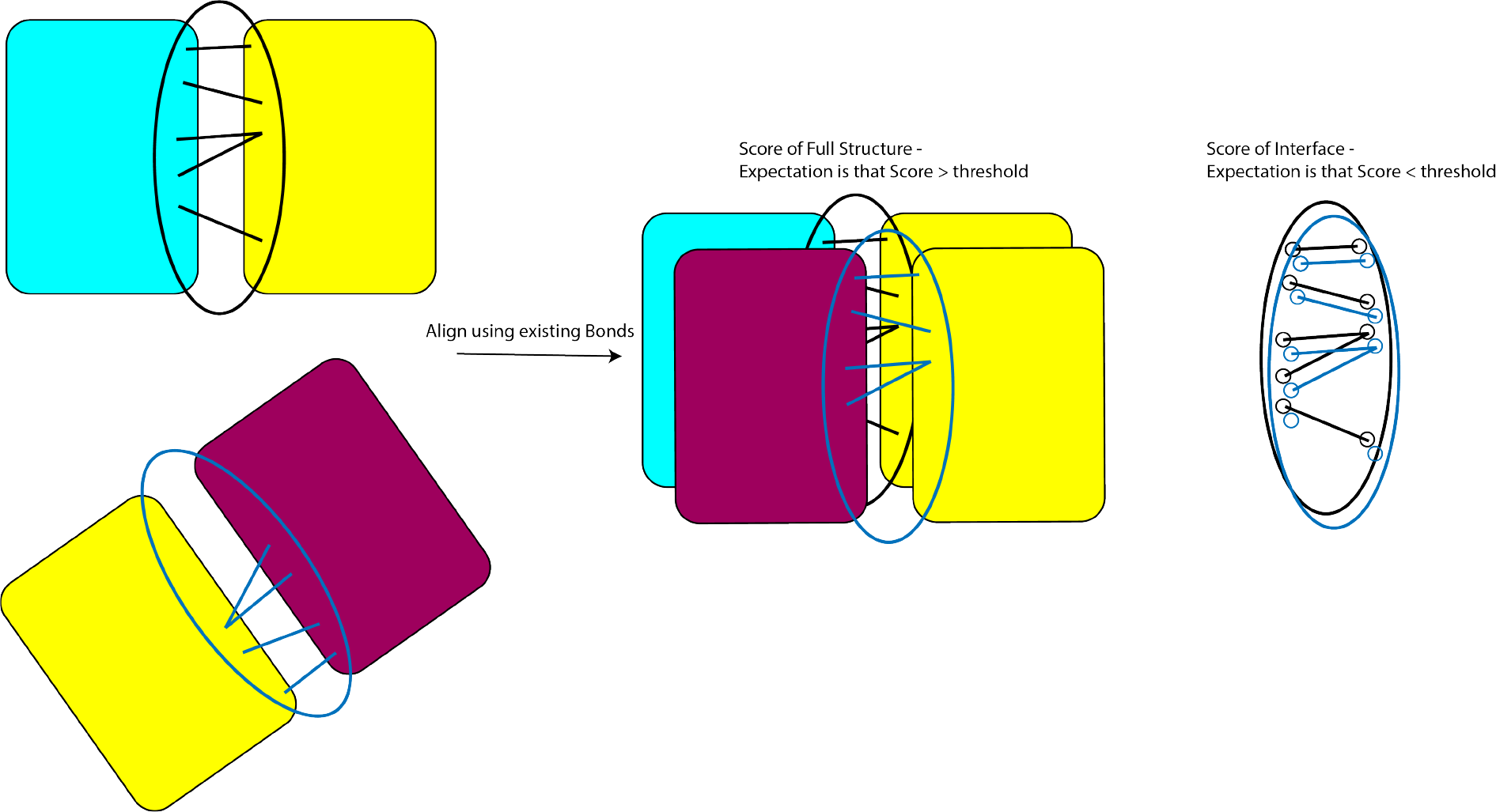
Methodology for obtaining a bond-anchored network structural alignment.

**Fig. 5.**
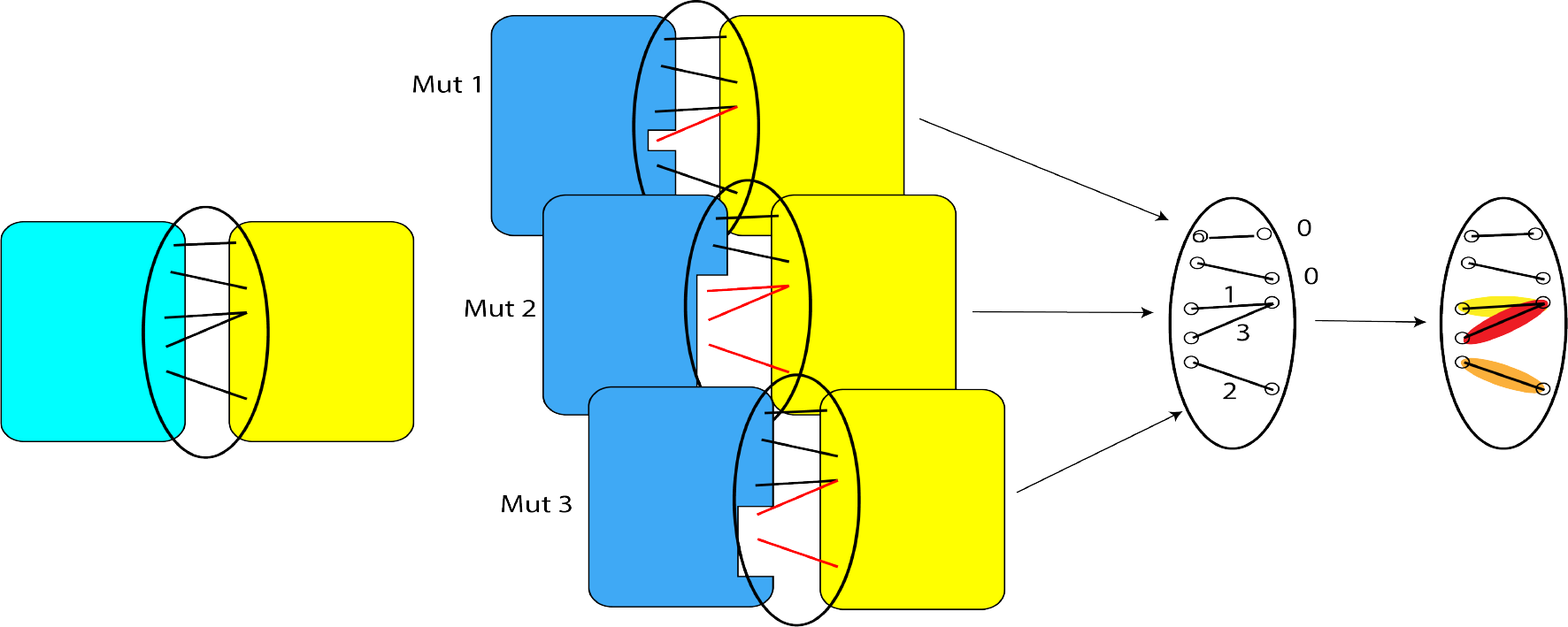
Methodology for producing a bond network heatmap from a set of mutations on a PPI.

**Fig. 6.**
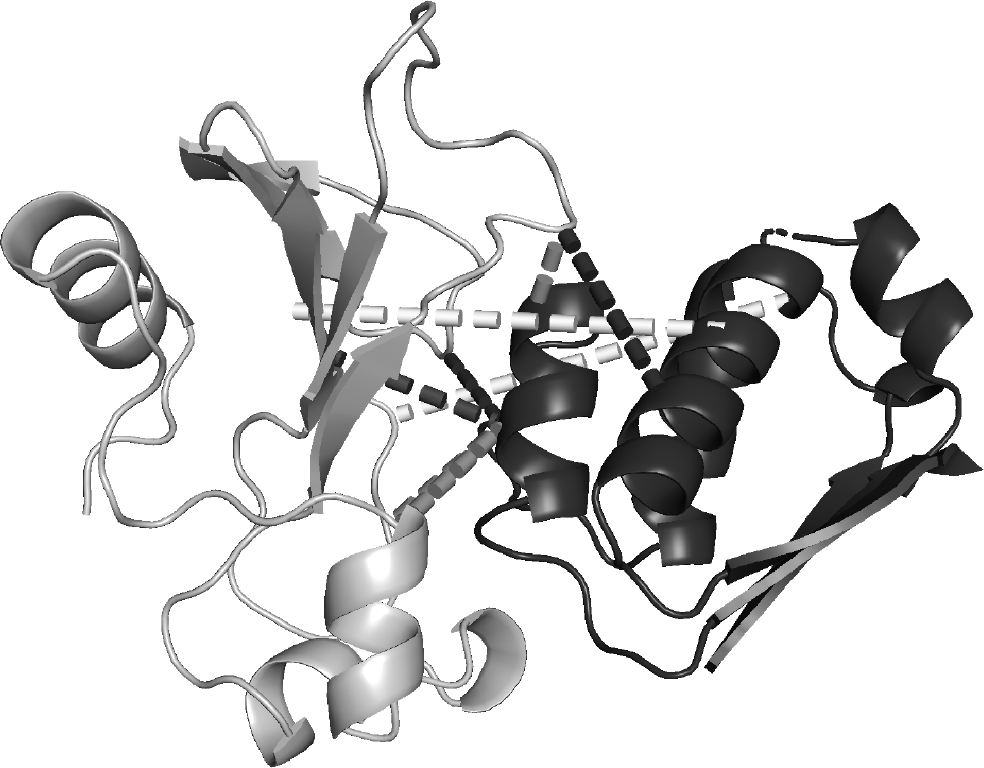
Bond network between barnase (white) and barstar (black) within a 3D model.

**Fig. 7.**
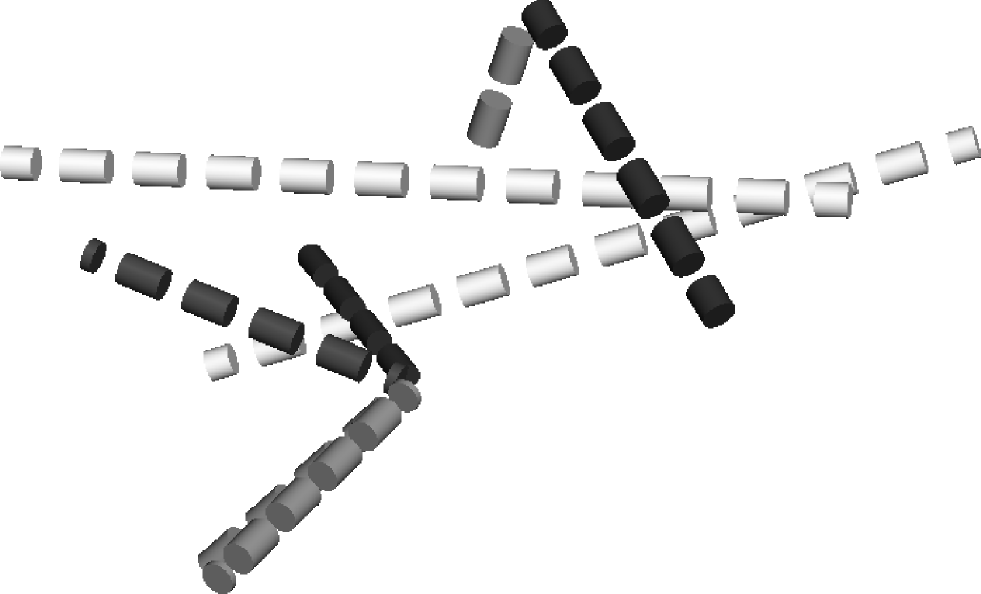
Visualization of a bond network in 3D.

**Fig. 8.**
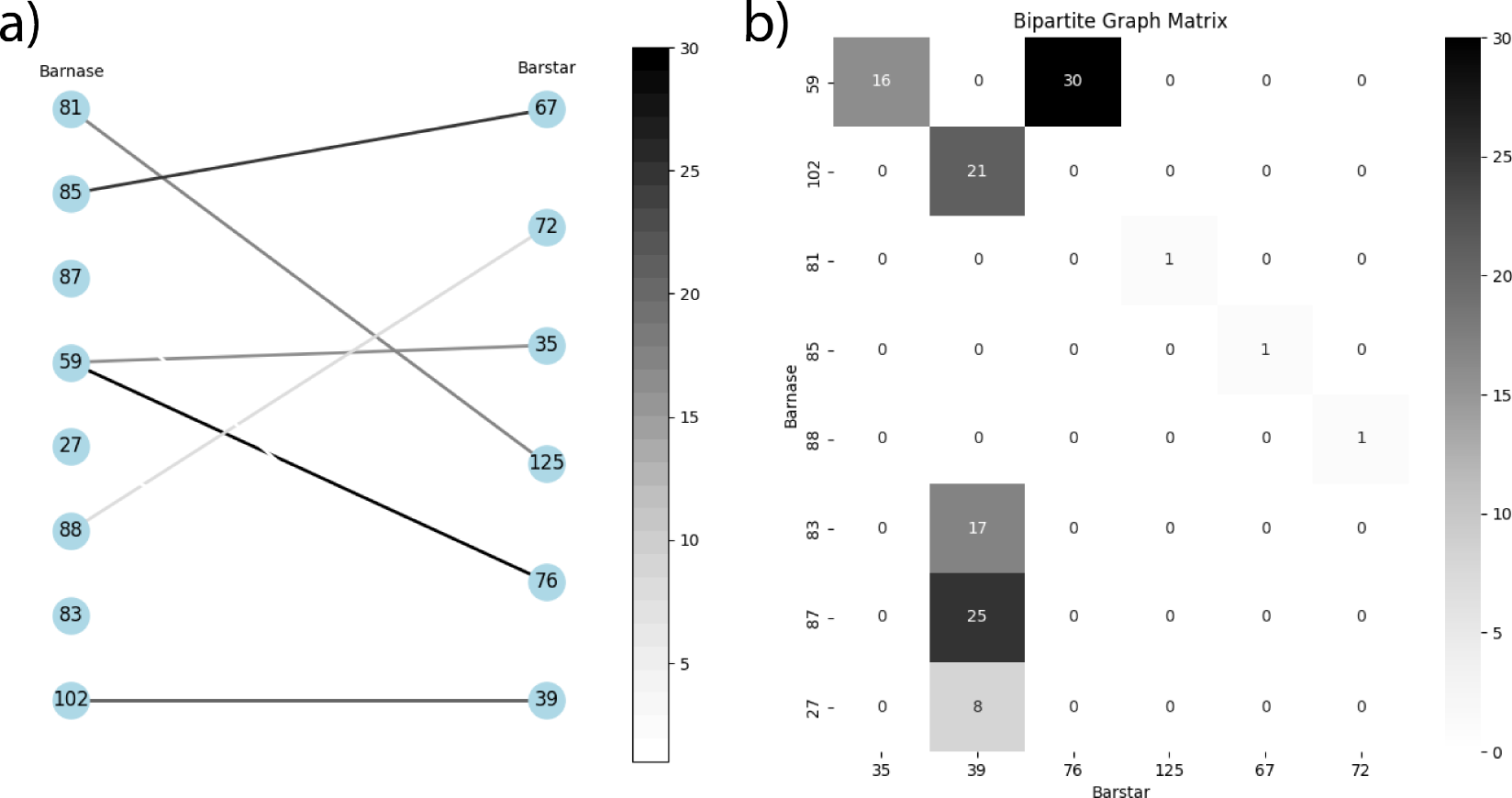
Data visualizations of heatmap data. a) Bipartite graph showing intermolecular connections from Barnase (left nodes) to Barstar (right nodes). Black edges indicate more changes to that bond compared to the wildtype due to mutation. b) A matrix representation of the graph. Greater total numbers across rows or columns indicate that bonds involving that node are less conserved due to mutation.

In order to determine the reliability of BANSA, we used multiple sources of validation techniques. A range of protein-protein interactions were investigated from the SKEMPI dataset for comparison of bond networks formed between interacting protein pairs. We aimed our analysis at establishing a correlation between the bond networks and the biochemical relevance of the PPI, to ensure the computational results are scientifically viable and biologically significant.

### A. Heatmap Analysis

One method for analyzing our bond networks was through the generation of heatmaps. In these visual representations, all individual bonds within a network were superimposed on each other using BANSA. This enabled a consolidated view of the entire bond network, allowing for intuitive and comprehensive interpretation of the intermolecular interactions within the proteins. The use of heatmaps helps us identify common and unique interaction patterns across different protein pairs, showing potential hotspots of interaction and regions of variability within the bond networks.

We show the heatmap results for barnase-barstar (1BRS) in, but similar results for serine proteases (1EAW, 3BN9) can be found in supplemental data. The heatmap of barnase-barstar ionic bonds show which bonds changed due to mutation as well as the number of times each bond changed from 74 different mutational experiments. In otherwords, we are superim-posing the difference between the wildtype bond network and each mutant bond network, represented as Σ|*G*_*wt*_ *− G*_*mut*_| for each bond that exists in the wildtype and mutant networks.

From a quick glance, we see that amino acids 59 and 102 on barnase have the most number of changes due to mutation (least conserved) with a total of 28 and 39 changes respectively. On barstar, amino acids 33 and 34 have 20 and 18 respectively. Cross-verification with barnase-barstar literature shows that barnase R59, H102 and barstar N33, N34 are all amino acids of significance and interest [1], [2].

### B. Literature Comparison

The final step in our validation process involved a thorough comparison of our findings against existing literature. The bond networks identified and the corresponding ΔΔG correlations were meticulously compared to documented protein-protein interactions in our DiffBond methods paper, Tam et al [3]. This cross-verification served to establish the accuracy and reliability of our method by aligning the computationally derived bond networks with experimentally validated interaction data.

Our comprehensive analysis, integrating heatmap visualizations, ΔΔG correlations, and comparison with literature, will attempt to validate our BANSA approach, confirming its capability to accurately represent and analyze protein-protein interactions through a graphical and bond-anchored representation of the interface.

## III Discussion

